# Environmental, Developmental, and Genetic Conditions Shaping Monarch Butterfly Migration Behavior

**DOI:** 10.1101/2025.01.16.633431

**Authors:** Hsiang-Yu Tsai, Cristian Molina, John Pleasants, Marcus R. Kronforst

## Abstract

Monarch butterflies in North America migrate south each autumn, but the mechanisms that initiate their migratory flight remain incompletely understood. We investigated environmental, developmental, and genetic factors that contribute to directional flight by testing summer and autumn-generation monarchs in three flight simulators: two at ground level (with and without wind blockage) and a novel balloon-based system that raised butterflies 30 meters into the air. Monarchs reared under autumn-like conditions in a growth chamber during the summer were also tested to explore the influence of developmental cues. Autumn generation monarchs demonstrated significant southwestern flight orientation, observed exclusively in the balloon simulator, underscoring the importance of high-altitude flight for migratory behavior. Summer generation monarchs reared under autumn-like conditions displayed southward orientation, larger wing sizes, and partial reproductive diapause, indicating specific seasonal environmental cues that are sufficient to induce migratory traits. In contrast, a lab line of monarchs reared in captivity since 2016 exhibited diminished wing size and reduced orientation ability, even when raised outdoors in the autumn, consistent with a loss of migratory traits in the absence of migration. Surprisingly, butterflies in the balloon simulator tended to orient upwind, which suggests that wind may also serves as a directional cue during migration. These findings highlight the critical roles of altitude, wind, and environmental cues in monarch migration and validate the balloon flight simulator as a powerful tool for studying migratory behavior. This research advances our understanding of the initiation of monarch migration and informs strategies for conservation efforts amidst environmental change.

## Introduction

Migration is a vital strategy used by many animal species to access seasonal resources or track favorable environmental conditions. This complex behavior depends on the timing of departure, which influences stopover patterns, arrival timing, and ultimately the fitness of migrating individuals (1–3). Understanding the triggers that initiate migration is therefore key to understanding how migrants adapt to their environments (4, 5). The onset of long-distance migration involves two interconnected components: a developmental phase that prepares the organism for migration (6–8) and a flight strategy that enables navigation through varying environmental conditions (9–11). During development, migrants undergo physiological, morphological (12), and metabolic changes (13) that equip them for sustained long-distance travel, with these traits shaped by genetic and environmental factors. In addition, effective flight strategies allow migrants to respond to dynamic environmental cues such as sun position (14), wind conditions (9, 15), and temperature (16, 17), ensuring successful navigation and energy efficiency during migration.

Among migratory species, insects are especially numerous, both in terms of biodiversity and ecological significance, serving as important pollinators and decomposers (18–20). Particularly, the monarch butterfly (*Danaus plexippus*) is a well-known migratory species that is famous for the long-distance migration of the eastern North American population. Unlike many migratory animals, which repeat their migration multiple times, eastern North American monarchs migrate only once in their lifetime. In response to seasonal cues, monarchs eclosing in the autumn embark on a southward journey of up to 4,000 kilometers to their overwintering sites in central Mexico. In spring, these same individuals fly north to the southern United States, where they reproduce and die, with successive generations continuing the northward journey (21–23). This multi-generational migration precludes the possibility of learning the migration route, indicating that this behavior is innate.

Migratory monarchs display distinctive traits, including southern-oriented flight in autumn, relatively large forewings, and reproductive diapause. Previous studies have explored the environmental cues influencing these traits (24–30), with particular attention placed on factors such as decreasing day length and temperature (30). Research has also delved into the neural mechanisms underlying migration (31, 32) and how monarchs utilize flight strategies to navigate environmental conditions, employing methods such as radio tracking (28, 33, 34) and vanishing bearings (35, 36). However, while considerable progress has been made in understanding the developmental processes that initiate migration, these studies have predominantly focused on the impact of rearing environments, as these are relatively straightforward to manipulate through common garden experiments. In contrast, comprehensive investigations into the interplay between developmental processes and flight strategies—especially how both factors combine to trigger migration—remain scarce due to the challenges of tracking small insects over long distances and the low likelihood of retrieving released butterflies.

In this study, we addressed two critical aspects of monarch butterfly migration: (1) how genetic background and environmental conditions during development shape the initiation of migration, and (2) how flight strategies—such as directional orientation, altitude preferences, and wind utilization—interact with environmental conditions to facilitate successful migration. To explore these questions, we reared monarch butterflies from summer and autumn generations under both controlled (growth chamber) and outdoor conditions, assessing migration-related traits under varying flight scenarios. Additionally, we tested monarchs from a long-term lab-reared population to evaluate how intermediate numbers of captive generations affect migratory behavior, particularly the retention or loss of directional orientation. Using a novel balloon-based flight simulator alongside ground-based systems, we examined the contributions of environmental and developmental cues to migratory traits.

Our findings emphasize the critical roles of altitude, wind, and seasonal cues in the initiation of migration. They also reveal how migratory behavior diminishes in lab-reared populations over generations. These results provide new insights into the drivers of monarch migration, highlighting the challenges faced by these butterflies and the importance of preserving the environmental conditions essential for their migratory success.

## Results and Discussion

To investigate the flight strategies that monarch butterflies use during the initiation of migration, we captured wild North American monarchs and reared them outdoors for both summer and autumn generations (hereafter, “summer field” and “autumn field”, Fig S1). We then tested their orientation behavior in the autumn using three different flight simulators that mimicked various flight conditions: (1) ground level open air, representing natural flight conditions, (2) ground level wind block, simulating a controlled environment at ground level, and (3) the balloon method, where butterflies experienced ascent to a 30-meter height with full wind exposure (Fig 1A). The ground-level simulators did not provide a height stimulus, while the transparent wind block allowed butterflies to perceive their surroundings visually but without wind interaction. In the balloon method, butterflies experienced both altitude and wind conditions similar to those in the wild, allowing us to simulate their migration flight more realistically.

**Figure 1.**
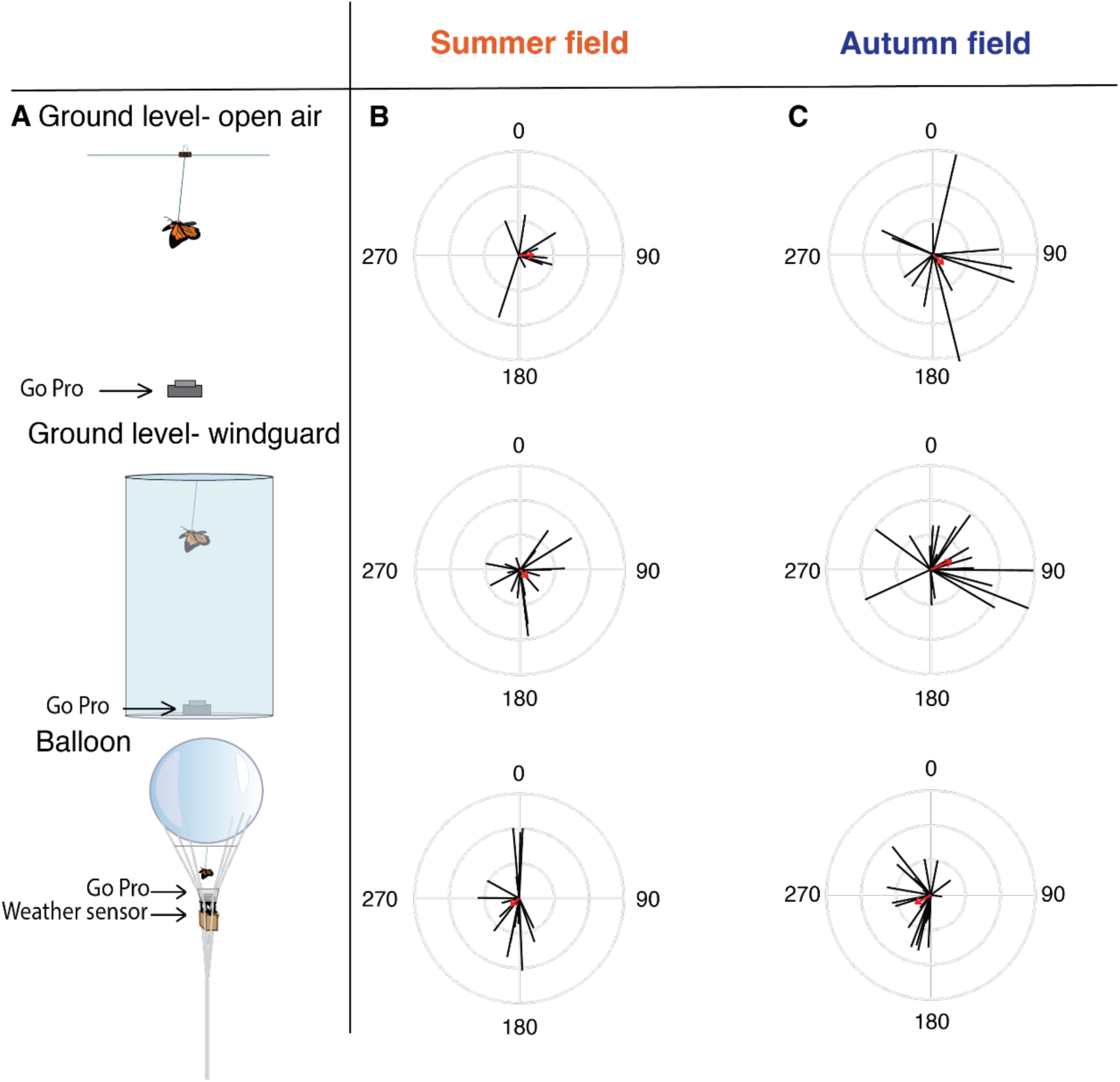
Monarch butterfly orientation behavior in field-reared populations across flight simulators. (A) Schematic diagram of the three flight simulators: two open-air ground-level simulators, one with a wind guard, and the balloon flight simulator. The GoPro camera beneath the butterfly records orientation behavior during flight. (B, C) Orientation plots for Eastern North American monarchs reared in the field, comparing the Summer (B, ground level open air: n = 10, Rayleigh test, P = 0.19; ground level with wind block: n = 19, Rayleigh test, P = 0.15; balloon: n = 18, Rayleigh test, P = 0.37) and Autumn generations (C, ground level open-air: n = 14, Rayleigh test, P = 0.4; ground level with wind block: n = 20, Rayleigh test, P = 0.017, balloon: n = 20, Rayleigh test, P = 0.017) across the three simulators. Each black line indicates the mean flight direction of an individual butterfly (0°–359°), with line length proportional to the strength of directional preference (0–1). The red arrow represents the mean group direction. Directions are plotted relative to 0° (north) and 180° (south). Autumn-generation butterflies oriented south only in the balloon flight simulator.

We recorded each butterfly’s average flight direction (0° to 359°) and the consistency of its flight (0 to 1) across all three simulators. In the summer field generation, butterflies exhibited no significant directional orientation across any flight simulator (ground level open air: σ = 86°, n = 10, r = 0.092, Rayleigh test, P = 0.19; ground level with wind block: σ = 132°, n = 19, r = 0.072, Rayleigh test, P = 0.15; balloon: σ = 230°, n = 18, r = 0.068, Rayleigh test, P = 0.37, Fig 1B), aligning with previous findings that monarchs do not migrate in summer and thus show no directed flight pattern (24, 37). In contrast, the autumn field generation, presumed to be migratory, showed strong southwest directionality exclusively in the balloon flight simulator (σ = 240°, n = 20, r = 0.11, Rayleigh test, P = 0.017, Fig 1C). Ground-level simulators, with and without wind blocking, yielded no significant southern orientation (open-air: σ = 134°, n = 14, r = 0.1, Rayleigh test, P = 0.4; wind block: σ = 67°, n = 20, r = 0.16, Rayleigh test, P = 0.017, Fig 1C). These results support existing evidence that high-altitude flight enables monarchs to integrate visual height cues and directional information from the sun, enhancing migratory orientation (10). Such behavior is analogous to findings in migratory birds, where high-altitude release after visual orientation results in sustained migratory behavior (38).

To evaluate environmental effects on orientation, we reared summer-generation butterflies in a growth chamber under autumn-like conditions (hereafter referred to as “autumn growth chamber”, Fig S1) and tested their orientation using the balloon simulator. Additionally, we tested butterflies from a lab population reared in captivity since 2016/2017, which we then raised outdoors under conditions similar to field populations (hereafter referred to as “summer lab line” and “autumn lab line”, Fig S1). Due to the high mortality rate of the summer lab line population, we were unable to test the orientation behavior of this group.

In the balloon test, butterflies from the autumn growth chamber treatment exhibited strong directional orientation (mean vector angle = 214°, n = 23, r = 0.17, Rayleigh test, P < 0.001; Fig. 2A), indicating that autumn-like developmental conditions effectively prime migratory behavior. This result has significant implications. While previous studies have struggled to pinpoint the specific developmental conditions that trigger migration in monarch butterflies (24, 27), our findings demonstrate that gradual decreases in day length and temperature, mimicking natural seasonal variation, are sufficient to induce southern flight orientation (39). This highlights the critical role of environmental cues in initiating migratory behavior.

**Figure 2.**
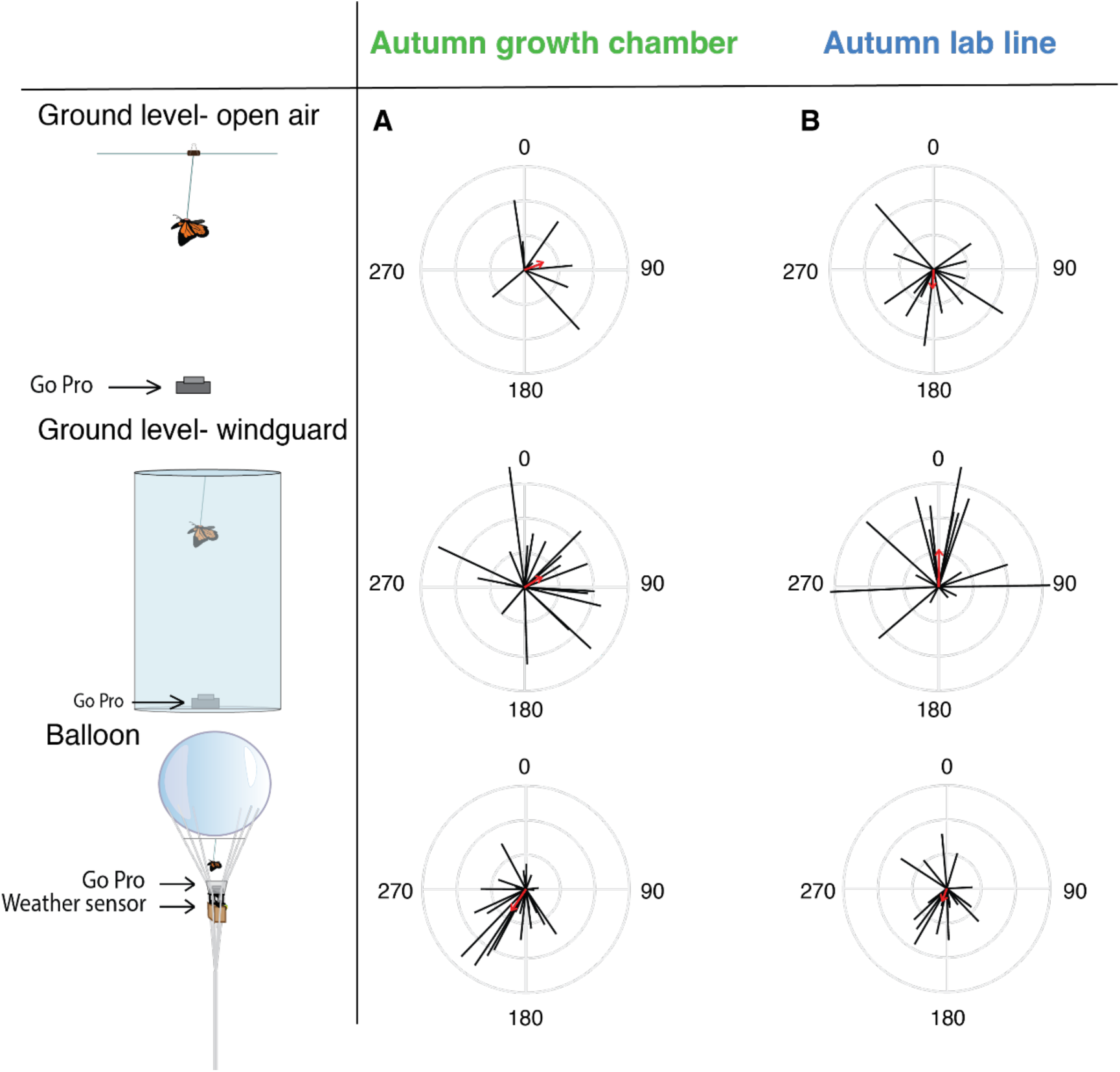
Orientation behavior of monarch butterflies reared in autumn-like conditions and the laboratory-maintained autumn generation. (A) Orientation plots of summer generation monarchs reared in a autumn-like growth chamber, tested across three flight simulators. Butterflies oriented south only in the balloon flight simulator, with no significant southern orientation in the ground-level simulators (ground level open-air: n = 8, Rayleigh test, P = 0.29; ground level with wind block: n = 21, Rayleigh test, P = 0.092; balloon: n = 23, Rayleigh test, P < 0.001). (B) Orientation plots of autumn lab line monarch butterflies, which showed only marginally significant southern orientation behavior in the balloon flight simulator (ground level open-air: n = 14, Rayleigh test, P = 0.18; ground level with wind block: n = 17, Rayleigh test, P = 0.008; balloon: n = 21, Rayleigh test, P = 0.047). Each black line represents the mean flight direction of an individual butterfly (0°–359°), with line length proportional to the strength of directional preference (0–1). The red arrow indicates the mean group direction. Orientation is plotted relative to 0° (north) and 180° (south).

For the autumn lab line, southern directional orientation in the balloon simulator was marginally significant (σ = 200°, n = 21, r = 0.09, Rayleigh test, P = 0.047, Fig 2B). Thus, evidence for southern orientation in this group is limited, likely due to prolonged captive rearing. This diminished orientation behavior, coupled with reduced immune function and survival rates observed in lab populations relative to field populations, may reflect a decline in migratory traits over time.

We further examined orientation behavior using ground-level simulators with autumn growth chamber and autumn lab line populations, both of which displayed no significant southern orientation (autumn growth chamber: ground level open-air: σ = 70°, n = 8, r = 0.14, Rayleigh test, P = 0.29; ground level with wind block: σ = 58°, n = 21, r = 0.14, Rayleigh test, P = 0.092, Fig 2A; autumn lab line: ground level open-air: σ = 183°, n = 14, r = 0.12, Rayleigh test, P = 0.18; ground level with wind block: σ = 1°, n = 17, r = 0.26, Rayleigh test, P = 0.008, Fig 2B). Consistent with our results with the autumn field generation, these results demonstrate that butterflies do not exhibit migratory orientation at ground level.

Although our study demonstrated that monarchs did not exhibit migratory orientation in the ground-level simulators we tested, it is important to acknowledge that prior research using the traditional fixed-tether monarch flight simulator—a ground-level device enclosed in an opaque cylinder—has shown southern orientation (40, 41). The traditional simulator differs in key ways from the ground-level simulators in our study, which lacked full enclosure and used a free tether design, potentially accounting for the differing results. Notably, the balloon flight simulator employed here had higher participation rates (almost 100%) compared to both the ground-level simulators and the traditional fixed-tether flight simulator, indicating its utility as an improved method for assessing orientation behavior. This enhanced participation, combined with the robust southern orientation observed in the balloon method, highlights its value for studying insect migratory behaviors under natural conditions.

Reproductive status and wing size, both migration-linked traits, were assessed after orientation behavioral tests for all groups (summer lab line, summer field, autumn growth chamber, autumn lab line, autumn field). All females from the summer lab line and summer field groups were reproductively mature (summer field: n = 28, summer lab line: n = 7, Fig. 3A), with a mean of 90 or more mature oocytes per female (summer field: n = 28, 93.4 ± 9.8 SE; summer lab line: n = 7, 148 ± 24 SE, Fig. 3B). In contrast, fewer than 20% of females from the autumn lab line and autumn field groups were reproductively mature (autumn lab line: n = 64, mature ratio = 0.17 ± 0.048 SE; autumn field: n = 41, mature ratio = 0.12 ± 0.052 SE, Fig. 3A). Additionally, mature oocyte counts were significantly lower in these groups than in summer butterflies (autumn lab line: n = 64, 5.56 ± 2 SE oocytes; autumn field: n = 41, 1.44 ± 0.76 SE, Mann–Whitney U test, P < 0.001, Fig 3B), consistent with previous observations that autumn generation, migratory monarchs are in reproductive diapause (25, 43).

**Figure 3.**
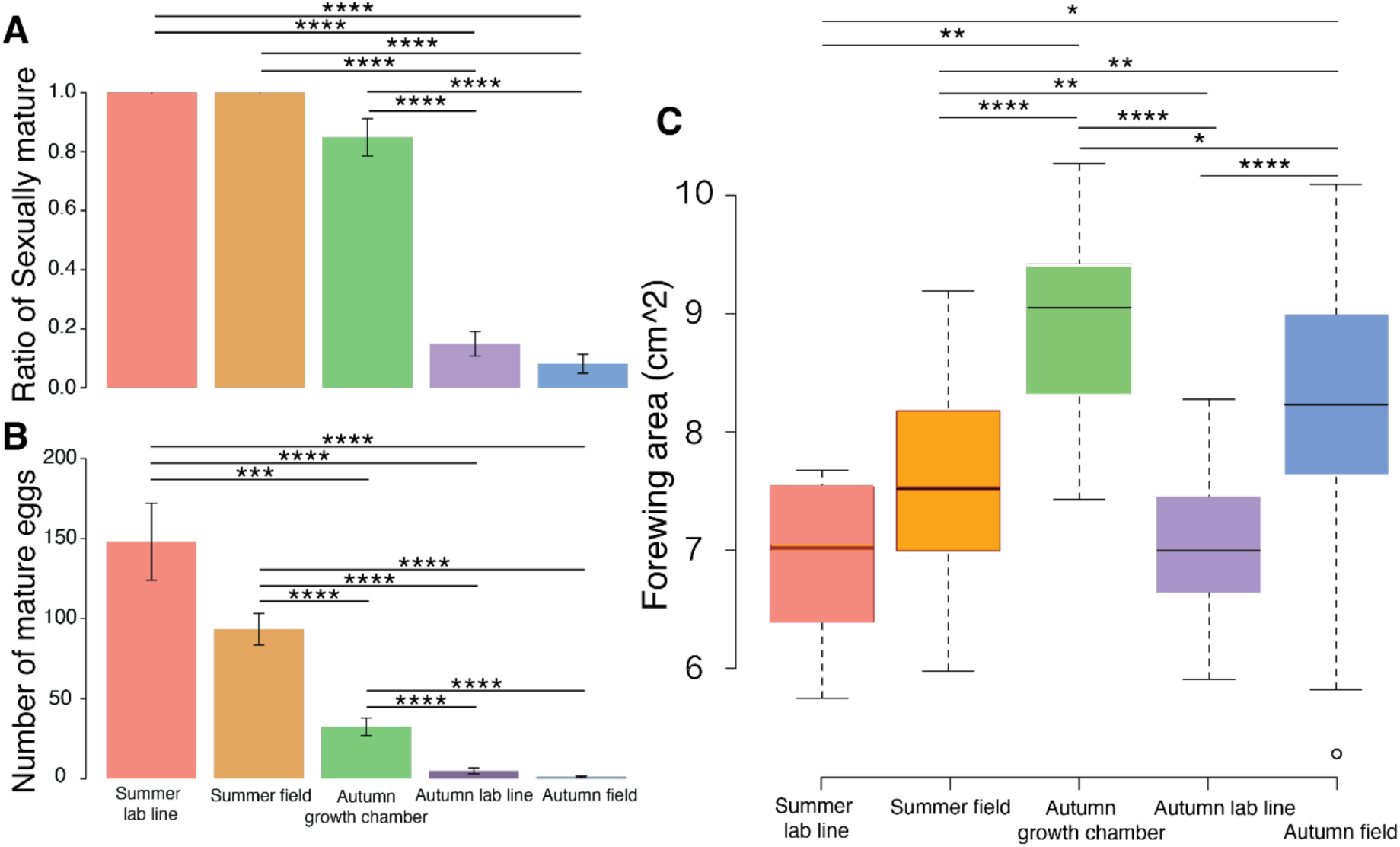
Reproductive status and forewing size differ among monarch butterfly treatment groups. (A) Proportion of sexually mature females (summer lab line: n=7; summer field: n = 28; autumn growth chamber: n=39; autumn lab line: n = 64; autumn field: n = 41). and (B) number of mature oocytes in the ovaries of females across five treatment groups: summer lab line, summer field, autumn growth chamber, autumn lab line, and autumn field generation (summer lab line: n=7; summer field: n = 28; autumn growth chamber: n=39; autumn lab line: n = 64; autumn field: n = 41). (C) Forewing area of butterflies across the five treatment groups, with autumn field and autumn growth chamber butterflies generally exhibiting larger forewing areas (summer lab line: n=7; summer field: n = 49; autumn growth chamber: n=26; autumn lab line: n = 114; autumn field: n = 72). The standard error was presented as the error bar. The number of asterisks indicates the level of statistical significance: *P < 0.05, **P < 0.01, ***P < 0.001, ****P < 0.0001.

Interestingly, butterflies reared in the autumn growth chamber treatment exhibited intermediate reproductive traits compared to summer and autumn field females. Although these butterflies had a higher reproductive maturity ratio than autumn field butterflies (autumn growth chamber: n = 39, mature ratio = 0.85 ± 0.059 SE; autumn field: n = 41, mature ratio = 0.12 ± 0.052 SE, Mann– Whitney U test, P < 0.001, Fig. 3A), they stored fewer oocytes than summer females (autumn growth chamber: n = 39, 31.3 ± 4.9 SE; summer field: n = 28, 93.4 ± 9.8 SE, Mann–Whitney U test, P < 0.001, Fig. 3B). This pattern suggests that reductions in day length and temperature can influence the induction of reproductive diapause. However, the differences between the autumn growth chamber and autumn field butterflies indicate that some aspects of the natural autumn environment that are not replicated in our growth chamber, such as exposure to the position of the sun (42), may contribute to reproductive diapause.

We next examined forewing size across all butterfly groups, as larger forewings are typically associated with migratory monarch populations, aiding in long-distance travel (43). Autumn-generation butterflies raised in the field had significantly larger forewing sizes compared to summer-generation individuals (summer field: n = 49, forewing area = 7.58 ± 0.12 cm²; autumn field: n = 72, forewing area = 8.27 ± 0.12 cm², Mann–Whitney U test, P < 0.001, Fig. 3C), consistent with their migratory behavior. Interestingly, butterflies from the summer generation raised in the autumn-like growth chamber exhibited even larger forewing areas than those from the autumn field generation (n = 26, forewing area = 8.97 ± 0.14 cm², Mann–Whitney U test, P < 0.001, Fig. 3C). However, the captive lab colony monarchs, regardless of season, displayed smaller forewing sizes compared to wild-derived individuals (summer lab line: n = 7, forewing area = 6.9 ± 0.3 cm², Mann–Whitney U test, P = 0.0013; autumn lab line: n = 114, forewing area = 7.04 ± 0.056 cm², Mann–Whitney U test, P < 0.001, Fig. 3C). These results suggest that both environmental and genetic factors influence forewing size, a key migration-related trait.

The reduced forewing size and orientation behavior observed in the lab-line butterflies are consistent with trends documented in commercially bred monarchs (24), which may have been raised in captivity for decades. Commercial populations show smaller wings than wild-derived monarchs and fail to orient south in the autumn, yet they retain phenotypic plasticity related to reproductive diapause. The similarities between these commercial monarchs and our lab line, which has been reared in captivity since 2016/2017, provide insight into the temporal dynamics of migration trait loss under artificial rearing conditions. Furthermore, these trends in captivity parallel the natural loss of migratory behavior in monarch populations around the globe that historically dispersed out of North America (44). Non-migratory populations often exhibit smaller wings while retaining seasonal plasticity in female reproduction (45, 46) (though their directional flight orientation behavior remains unknown). These parallels suggest that evolution in long-term lab and commercial populations may reflect, to some extent, the dynamics of natural migration loss in monarchs. Our results demonstrate that migration-related traits such as orientation behavior and wing size can evolve very rapidly after a population transitions from migratory to sedentary.

Given that two of our flight simulators were open-air, we examined how butterflies interacted with wind during their flights. Fine-scale wind data were recorded during each experiment, though a sensor failure limited data collection for the ground-level open-air simulator. To assess the relationship between flight and wind, we compared the difference between each butterfly’s flight direction and wind direction. A smaller difference indicated alignment with the wind, while a larger difference indicated upwind flight. Among the autumn growth chamber butterflies tested in the ground-level simulator, the direction differences were uniformly distributed, indicating no significant relationship between flight orientation and wind (skewness = 0.077, n = 8, KS test, P = 0.55; Fig. 4A). In contrast, in the balloon-based simulators, the distributions of direction differences were negatively skewed across all treatment groups. This pattern revealed that most butterflies orienting southward in the balloon simulators flew against the wind (autumn lab line: skewness = −0.561, n = 21, KS test, P < 0.001; autumn field generation: skewness = −0.733, n = 20, KS test, P < 0.001; autumn growth chamber: skewness = −1.53, n = 23, KS test, P < 0.001; Fig. 4B).

**Figure 4.**
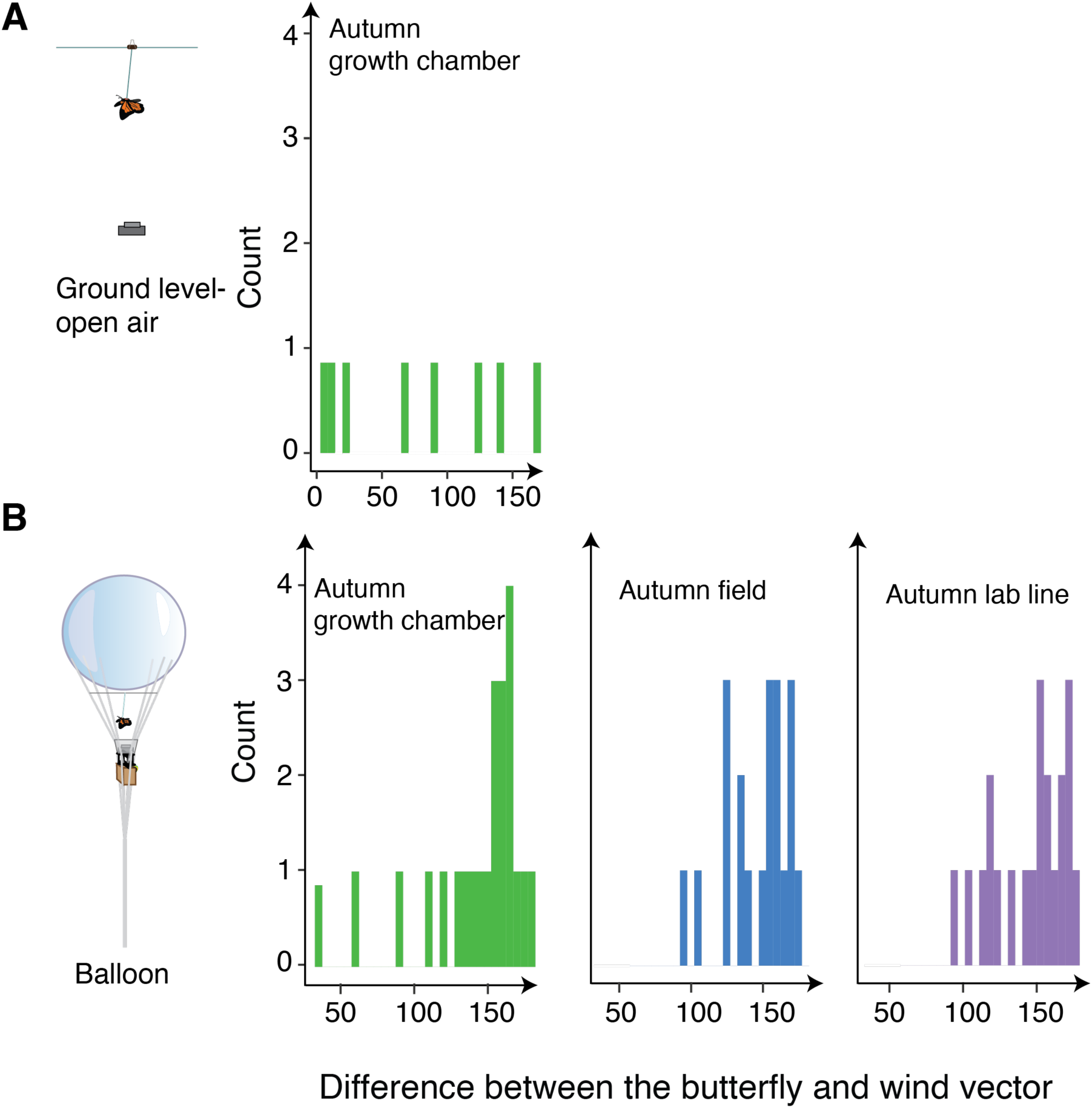
Interaction of migratory butterflies with wind across flight simulators. (A) In the open-air flight simulator at ground level, migratory butterflies showed no significant interaction with wind direction. (n = 8, P = 0.55). (B) In the balloon flight simulator, migratory butterflies consistently oriented upwind, flying against the wind (autumn growth chamber: n = 23, P < 0.001; autumn field generation: n = 20, P < 0.001; autumn lab line: n = 21, P < 0.001). The interaction between butterfly and wind direction was quantified as the difference between the butterfly’s mean heading and the wind direction during flight tests. Larger differences indicate more pronounced upwind flight, while smaller differences suggest alignment with the wind (tailwind).

Upwind flight is unusual for migratory animals, particularly monarch butterflies, which are generally thought to rely on tailwinds for long-distance travel (34). Furthermore, we observed no significant difference in wind interaction patterns between non-migratory summer field and migratory autumn field generations (summer field: n = 18; autumn field: n = 20, two-sample t-test, P = 0.42, Fig. S2). However, when wind direction and speed were explicitly incorporated into the analysis of butterfly flights in the balloon simulator, no significant directional wind pattern was detected for either the summer or autumn field generations (summer field: σ = 342°, n = 18, r = 0.46, Rayleigh test, P = 0.22; autumn field: σ = 0.5°, n = 20, r = 0.49, Rayleigh test, P = 0.16; Fig. S3A). These results suggest that monarchs in the balloon simulator actively selected their flight direction independent of wind conditions, reinforcing the idea that their orientation behavior is not dictated by wind direction.

In contrast, significant northward wind patterns were recorded during experiments with the autumn growth chamber and autumn lab line butterflies (autumn growth chamber: σ = 2°, n = 23, r = 1.1, Rayleigh test, P < 0.001; autumn lab line: σ = 341°, n = 21, r = 0.72, Rayleigh test, P = 0.0028; Fig. S3B). Long-term wind data from the European Centre for Medium-Range Weather Forecasts (ECMWF) corroborated these findings, revealing that prevailing winds in the Midwest during September—the peak migration period—blow predominantly from the south (Fig. S4). This aligns with previous studies reporting northward winds at migration starting points in southern Canada and the northern United States (47). These data suggest that migratory monarchs, like those in our balloon experiments, may initially fly upwind to orient themselves or to initiate migration. Such upwind behavior could serve as a mechanism for navigation, with prevailing wind patterns acting as important directional cues during the early stages of migration.

These findings raise intriguing questions about how monarchs detect and eventually exploit favorable winds to sustain their southward journey. Understanding the mechanisms by which migratory butterflies adjust their flight strategies in response to wind direction and strength could yield valuable insights into their navigation and energy conservation strategies. Further research is needed to explore how monarchs compensate for adverse winds and maintain their migratory trajectory en route to overwintering sites.

In summary, our findings highlight the complex interplay between environmental cues, physiological traits, and flight strategies in shaping monarch butterfly migration. By employing innovative flight simulator methods, we demonstrated that high-altitude flight conditions elicit strong migratory orientation, suggesting a critical role for visual and wind-related cues at these altitudes. Our results also underscore the influence of developmental conditions on migratory behavior and associated traits, such as forewing size and reproductive diapause, while emphasizing the rapidity with which migratory characteristics are lost in sedentary populations. These insights not only advance our understanding of monarch migration but also provide a framework for investigating broader questions about the ecological and evolutionary factors driving animal migration (48). Future research should focus on unraveling the mechanisms underlying wind cue detection, the physiological adaptations enabling energy-efficient flight, and the potential genetic basis of migration-related traits. Such efforts will be pivotal for informing conservation strategies as migratory species face increasing environmental challenges (5).

## Materials and Methods

### Animal Husbandry

We collected monarch butterfly caterpillars from the field near Chicago in early June 2023 and housed them in 30.5 cm³ mesh pop-up cages with access to their host plant, *Asclepias syriaca* (common milkweed). To prevent *Ophryocystis elektroscirrha* (OE) infection, we collected fresh milkweed daily and cleaned the plants with a 1:19 bleach solution. Once adults eclosed, we screened each individual for OE spores using adhesive tape on their abdomens, and any infected butterflies were immediately frozen. OE-free butterflies were housed in medium-sized (91.5 cm × 30.5 cm²) mesh cages, separated by sex. This group constituted the “wild-caught” generation.

To produce the first captive-bred generation, we hand-paired males and females from the wild-caught generation. Mated females were placed individually in medium-sized cages with a host plant for egg-laying. Half of the eggs from each family were placed in an environmental chamber that simulated autumn conditions, including decreasing day length and temperature (see details below), to produce the “autumn growth chamber” treatment. The remaining eggs were reared outdoors at a garden on the University of Chicago campus, under natural summer conditions, generating the “summer field generation.”

Butterflies from the outdoor-reared summer generation were then hand-paired to produce the “autumn field generation,” which was reared outdoors at the University of Chicago’s Warren Woods Ecological Field Station. Throughout the process, all adult butterflies were individually labeled and fed Birds Choice butterfly nectar twice daily.

### Autumn**-**like Environmental Chamber

To replicate autumn-like environmental conditions, we recorded daily temperatures at the field station from August 5 to October 1, 2022, using a temperature sensor. Daytime and nighttime temperatures were calculated as averages for their respective periods. These data informed the settings of a growth chamber from July 5 to August 31, 2023, which simulated the recorded temperatures and day lengths from the 2022 field measurements. After October 1, 2022, the average temperature dropped below 10 °C, a threshold that could negatively affect butterfly survival or behavior. To address this, starting September 1, 2023, we adjusted the chamber to maintain autumn-like conditions conducive to butterfly survival. The adjusted daytime and nighttime temperatures were set to 20.5 °C and 15.7 °C, respectively, reflecting the September 2022 monthly averages, while day length continued to decrease incrementally to mimic the natural seasonal transition. The growth chamber used dual fluorescent lamps with fixed on/off timing that matched the programmed light schedule. These settings provided a controlled environment closely approximating the autumn conditions monarchs experience in the wild.

### Experiments with monarch lab lines

We conducted experiments using two separate lab colonies of monarch butterflies that have been maintained at Iowa State University since 2016 and 2017, respectively. From 2016/2017 to 2021, both colonies were reared in growth chambers set at 26.6°C with a 16-hour photoperiod. After 2021, the colonies were moved to classrooms without windows for educational purposes, where the butterflies were reared at ambient room temperature and artificial light conditions (15h dark: 9 h light). In July 2023, we received batches of eggs from both colonies, which were raised outdoors as separate families at our experimental location in Chicago. We used the same rearing protocol as for wild-caught caterpillars, feeding the caterpillars freshly cut and bleached *Asclepias syriaca* (common milkweed) daily. The adults from these caterpillars were combined to form the “summer lab line” experimental population.

Unfortunately, many individuals from the summer lab line died before flight tests could be conducted, preventing us from collecting orientation behavior data from this group. A small number of surviving adults were paired to generate an autumn generation, which was supplemented by a second batch of eggs from the 2016 and 2017 colonies, received in August 2023. The adults that emerged from this second round of caterpillars were combined with offspring from the summer lab line generation to form the “autumn lab line” population.

### Flight Simulator Methods

We used three different flight simulators to assess the directional flight orientation of individual butterflies under varying wind and altitude conditions. To ensure secure tethering, we first removed hair from the thorax of each butterfly using transparent tape, then applied a bandage to make the surface rougher for better attachment (49). A loose fishing line, attached to the bandage with UV glue, provided stability and allowed for more natural flight while also increasing survival rates. Butterflies were kept in mesh cages for at least one day to recover from the tethering process. Prior to testing, all butterflies were placed outdoors for at least one hour.

The first simulator method, the “ground level open air,” involved tethering the butterfly with a fishing line to a horizontal rope aligned with true north in an open area without any wind obstruction. The second method, the “ground level wind block,” was similar but included a large, 30 cm diameter transparent PVC cylinder to block wind, ensuring that the butterfly could fly without wind interference. A GoPro camera placed on a table directly beneath the butterfly recorded flight duration and direction for both methods.

For the “balloon method,” each butterfly was tethered with a fishing line beneath a 2.1-meter balloon, which was released to a height of 30 meters. This setup allowed the butterfly to fly at a high altitude while interacting with natural wind conditions (50). A GoPro camera and an LI-550F TriSonica weather sensor, positioned below the butterfly, recorded flight behavior alongside environmental variables, including wind speed, direction, temperature, and air pressure. A Raspberry Pi 4 connected to the weather sensor served as a data logger for weather metrics. Each test lasted 10 minutes and 30 seconds; however, if a butterfly stopped flying for more than 15 seconds, the test was terminated. To ensure accurate analysis, the first 30 seconds of each video were excluded, allowing the butterfly time to ascend and stabilize its flight. All experiments were conducted on sunny days between early September and October 2023 at the Warren Woods Ecological Field Station, a well-preserved natural site with minimal human disturbance.

### Flight Orientation Data Analysis

We used DeepLabCut version 2.2, a ResNET-50-based network, to train a tracking model to analyze butterfly heading directions. Five body parts—head, thorax, right forewing, left forewing, and abdomen tip—were labeled in the training videos. The output of the DeepLabCut model provided the coordinates of the five labeled body parts for each frame. To calculate the butterfly’s heading direction per second during the balloon flight simulator trials, we sampled every 60th frame, corresponding to the 60 fps GoPro framerate. For the two ground-level flight simulators, we sampled every 60th or 30th frame depending on the GoPro’s framerate. Only the head and thorax coordinates were used to determine the butterfly’s heading direction by subtracting the thorax coordinates from the head coordinates to create the flight vector.

For the two ground-level flight simulators, the butterfly was tethered to a white line oriented toward true north, which served as the reference for the north vector. The butterfly’s heading direction was determined by measuring the angle between its flight vector and the north vector. In the balloon flight simulator, heading direction was recorded using the LI-550F TriSonica weather station attached to the balloon, as the balloon could rotate while floating in the sky. A GoPro camera, aligned with the weather station’s reference direction, captured the butterfly’s flight vector. To calculate the butterfly’s heading direction, we first measured the angle between its flight vector and the long axis of the GoPro’s field of view (represented as the vector (0,1)). This angle was then added to the heading direction recorded by the TriSonica at the corresponding time frame, yielding the butterfly’s final heading direction.

To calculate the average heading direction for each trial, the butterfly’s heading was recorded every 10 seconds. These angular data (in degrees) were converted into Cartesian coordinates using the “Circular” package in RStudio 1.3.959. The mean heading direction (0–359°) and flight consistency (r = 0–1, where higher values indicate greater consistency) were then computed for each butterfly. To determine whether butterflies within each group exhibited significant directional flight, we weighted each individual’s mean flight direction by its consistency score and conducted a Rayleigh test using the “astropy.stats.rayleightest” package in Python.

### Ovary and Wing Analyses

To assess reproductive status, we froze all butterflies post-experiment and dissected female ovaries. The abdomens of the female butterflies were removed, and a longitudinal cut was made to open the abdomen. We classified an oocyte as mature if it had surface ridges typical of those about to be laid. We counted the number of mature oocytes inside the abdomen, and individuals with more than one mature oocyte were considered sexually mature. Those without were classified as being in reproductive diapause.

After flight tests, we removed and analyzed the right forewing of each butterfly to measure its area. Photographs of the wings were taken, and the images were converted to binary form. The software ImageJ was then used to calculate wing area.

We compared physiological and morphological traits across different generations and treatment groups using the Mann-Whitney U test. For reproductive traits, we assessed binary data indicating whether individuals had reached maturity as well as the number of mature oocytes. Wing variables were also analyzed using the Mann-Whitney U test to evaluate differences across groups.

### Wind Data Extraction

Real-time wind data were recorded during each flight experiment using two instruments: a TriSonica weather station mounted on the balloon and a HOBO by Onset S-WCG-M003 Ultrasonic Wind Smart Sensor at ground level. Both sensors collected wind speed and direction at one-second intervals throughout the flight tests. To analyze the butterflies’ flight behavior in relation to wind, we calculated the difference between the wind direction and the butterflies’ heading direction on a per-second basis. The distribution of these differences was plotted, and skewness was calculated to assess wind alignment. A negatively skewed distribution (skewness < 0) indicated upwind flight, while a positively skewed distribution (skewness > 0) suggested downwind flight. To evaluate whether the data distribution deviated from uniformity, we applied the Kolmogorov-Smirnov (KS) test. Additionally, to determine whether wind conditions during each test were significantly directional, wind direction was weighted by wind speed and analyzed using the Rayleigh test, following the same method applied to the butterflies’ flight direction.

We also obtained long-term wind data using the ERA5-Land reanalysis dataset from the European Centre for Medium-Range Weather Forecasts (ECMWF). To represent wind conditions across the Midwest, we selected six locations: the university garden (Lat: 41.80, Long: −87.59) and the field station (Lat: 41.84, Long: −86.62) where monarch butterflies were reared, along with four additional locations reported in the JourneyNorth program for monarch migration observations (Lat: 43.77, Long: −86.44; Lat: 41.57, Long: −88.49; Lat: 42.66, Long: −88.6; Lat: 41.56, Long: −87.51). Hourly wind speed and direction data for the month of September were extracted for each of these locations.

## Acknowledgments

We thank Sheng-Feng Shen for his help designing the balloon flight simulator, Matthias Steinrücken for his assistance with flight data analysis, Vaibhhav Sinha for his help coding on Raspberry Pi, and Keith Bidne for maintaining monarch lab colonies at Iowa State University and providing eggs. We also thank the Molina family for their support with transportation during fieldwork, the staff of the University of Chicago Warren Woods Ecological Field Station for their hospitality, and Trevor Price for comments on the manuscript. This research was supported by NSF grant IOS-1922624 and NIH grant R35 GM131828 to MRK.

## Figures and Tables

**Supplementary Figure 1.**
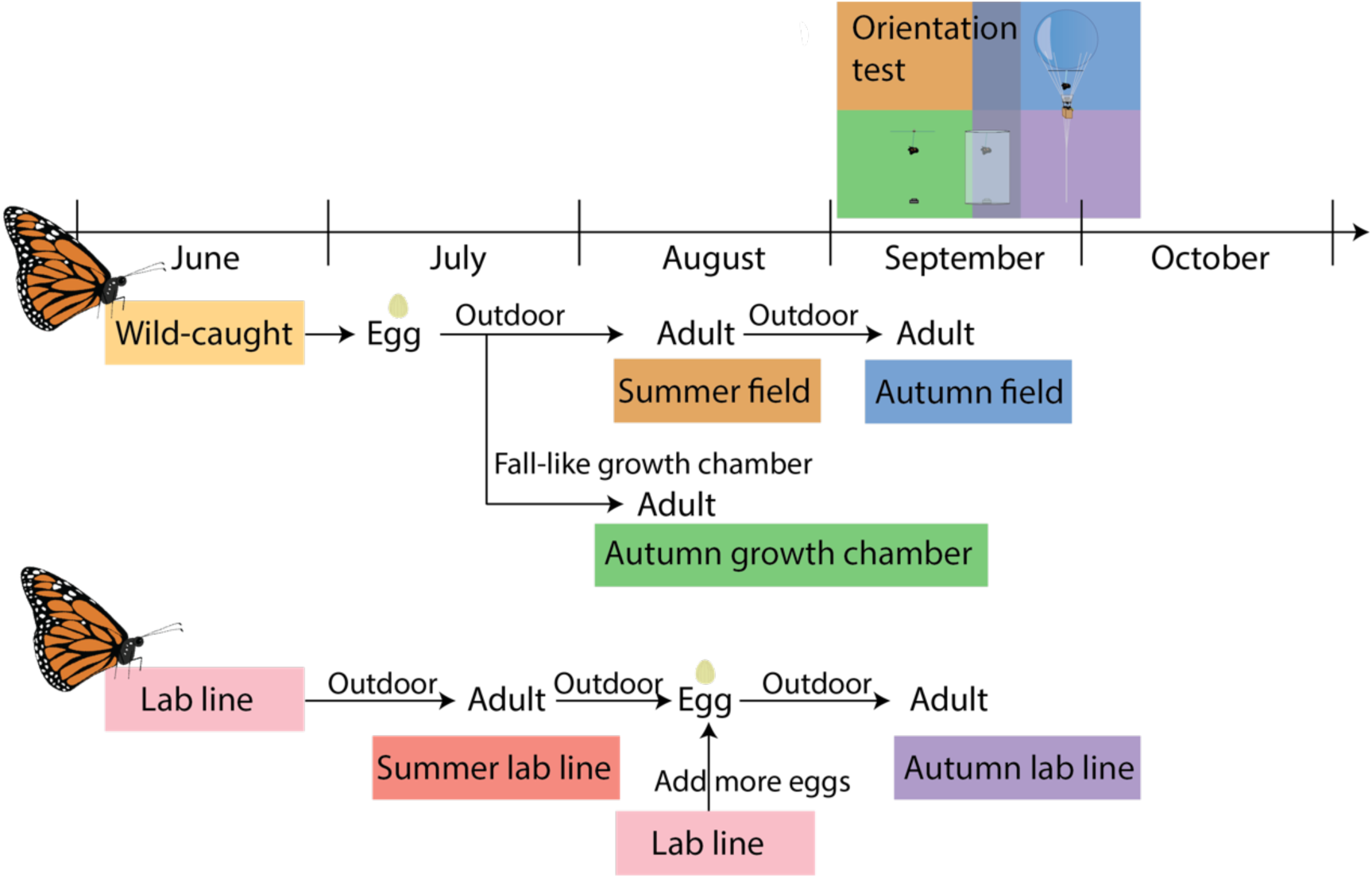
Schematic of rearing protocols and flight experiment timeline. Monarch butterflies were captured in the field in June and designated as the “wild-caught” generation. Eggs laid by wild-caught butterflies were split into two rearing conditions. Half were raised in an autumn-like growth chamber, and the resulting adults were classified as the “autumn growth chamber” group. The other half were reared outdoors, producing adults classified as the “summer field” group. Eggs laid by the summer field generation were reared outdoors, and their eclosed adults were classified as the “autumn field” group. Separately, monarch eggs obtained from a laboratory stock maintained at the Iowa State University since 2016/2017 were reared outdoors, with the resulting adults classified as the “summer lab line” group. Eggs laid by the summer lab line adults, along with a second batch of laboratory eggs, were also reared outdoors, producing adults classified as the “autumn lab line” group. Flight experiments were conducted on adults from the summer field and autumn growth chamber groups from early to mid-September, while adults from the autumn field and autumn lab line groups were tested from mid-September to early October.

**Supplementary Figure 2.**
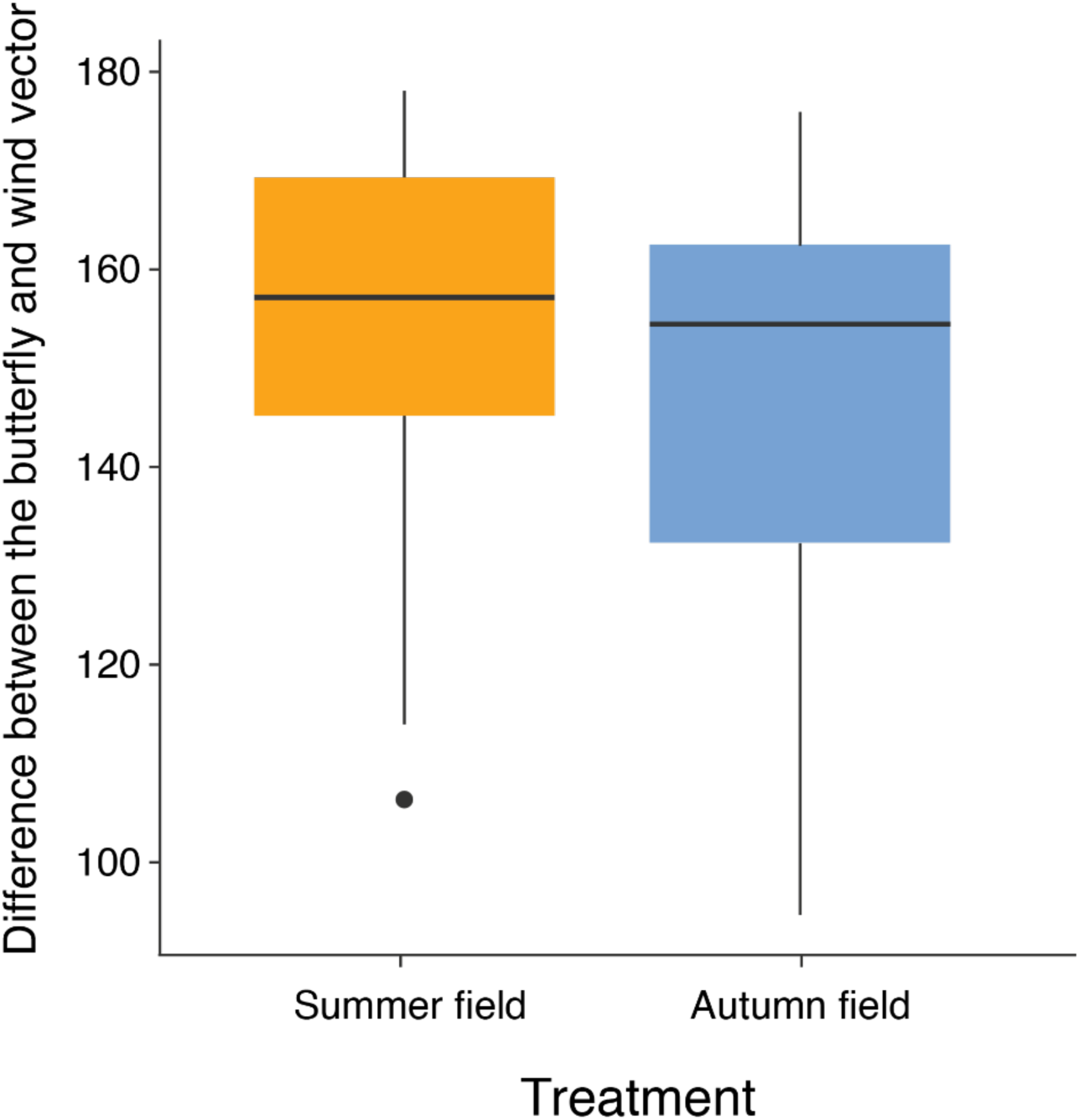
Comparison of wind interaction between Summer field and Autumn field groups. The difference between wind direction and butterfly flight direction was not significantly different between the summer field group (n = 18) and the autumn field group (n = 20) (two-sample t-test, P = 0.42). Error bars represent standard error.

**Supplementary Figure 3.**
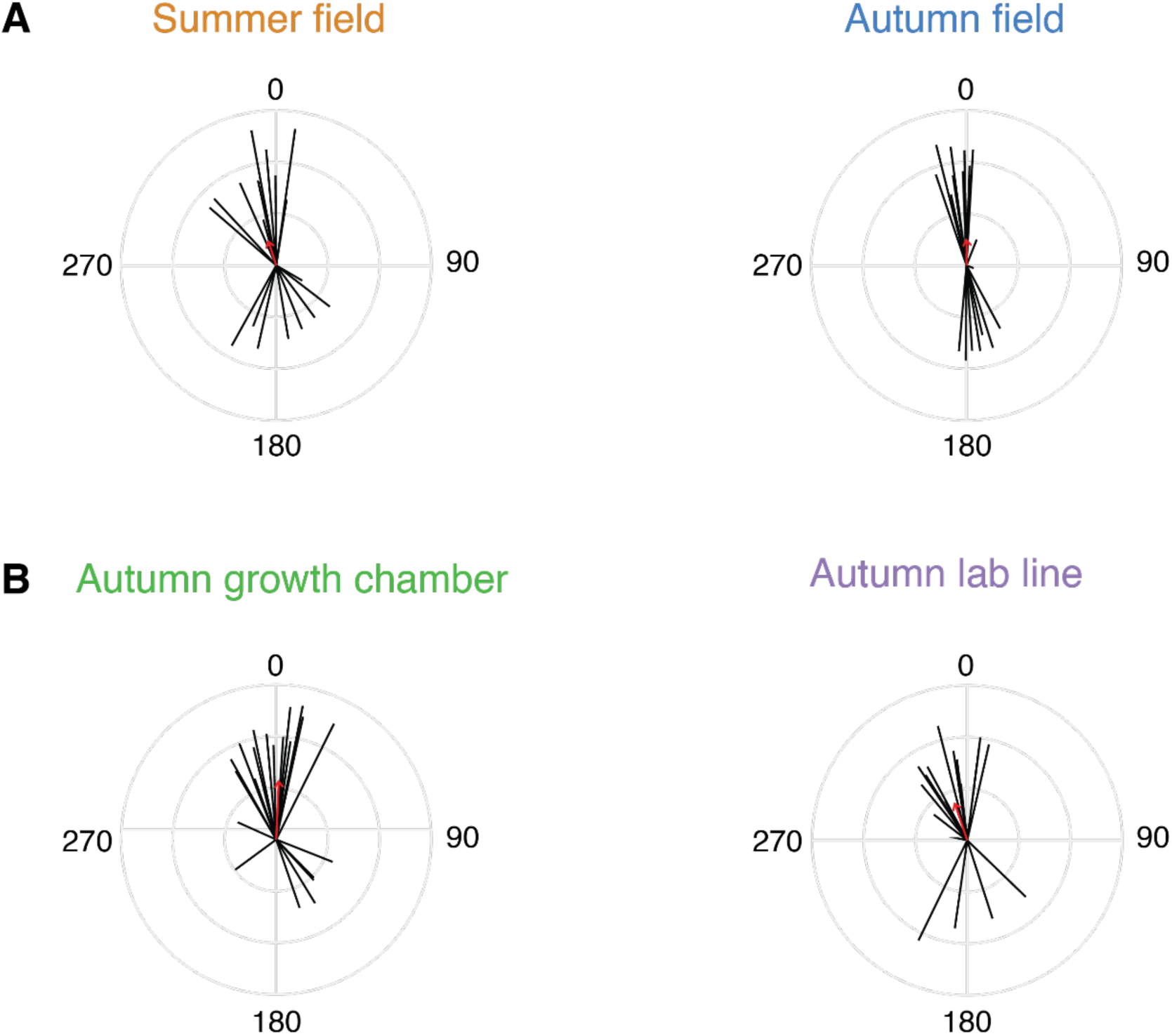
Variability in wind patterns during flight tests across treatment groups. Circular plots illustrate wind conditions experienced by butterflies during flight tests for (A) summer and autumn field generation groups (summer field: n = 18, Rayleigh test, P = 0.22; autumn field: n = 20, Rayleigh test, P = 0.16). and (B) autumn growth chamber and autumn lab line groups (autumn growth chamber: n = 23, Rayleigh test, P < 0.001; autumn lab line: n = 21, Rayleigh test, P = 0.0028). Each black line represents the mean wind direction for an individual test (0° to 359°), with line length indicating wind speed. The red arrow denotes the overall mean wind direction across tests for each treatment. 0° represents north, and 180° represents south.

**Supplementary Figure 4.**
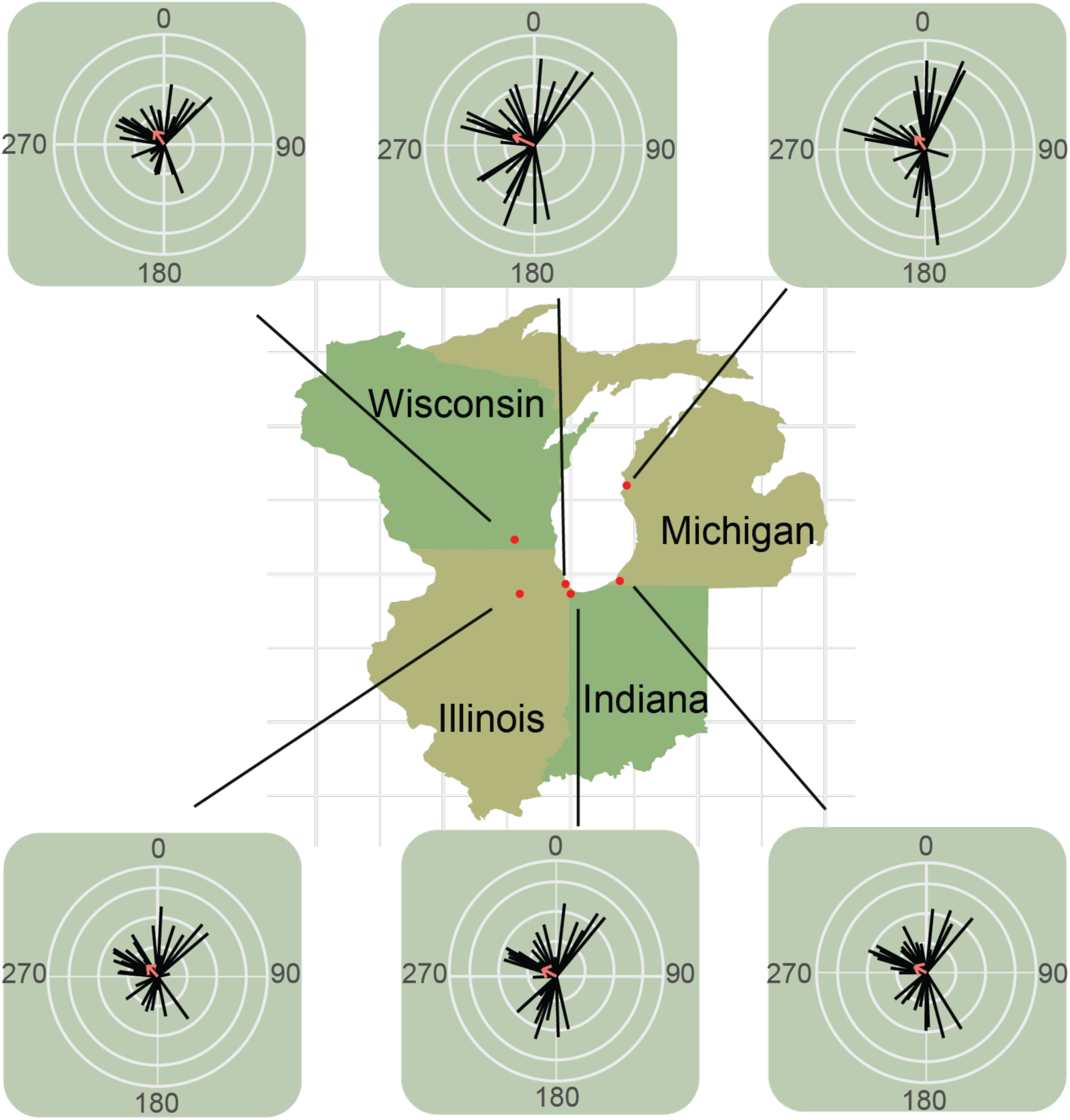
Prevailing wind patterns in the Northern Midwest during September. Circular plots show daily average wind direction and speed across six locations in September. Each black line represents the mean wind direction for a single day (0° to 359°), with line length corresponding to wind speed. The red arrow indicates the overall mean wind direction for the month at each location. Wind predominantly blew to the north during this period.

## Notes

https://doi.org/10.5281/zenodo.14668513

### Competing Interest Statement

The authors have declared no competing interest.

