## Supplementary figures and images for "Environmental, Developmental, and Genetic Conditions Shaping Monarch Butterfly Migration Behavior"

### Supplementary Figure 1

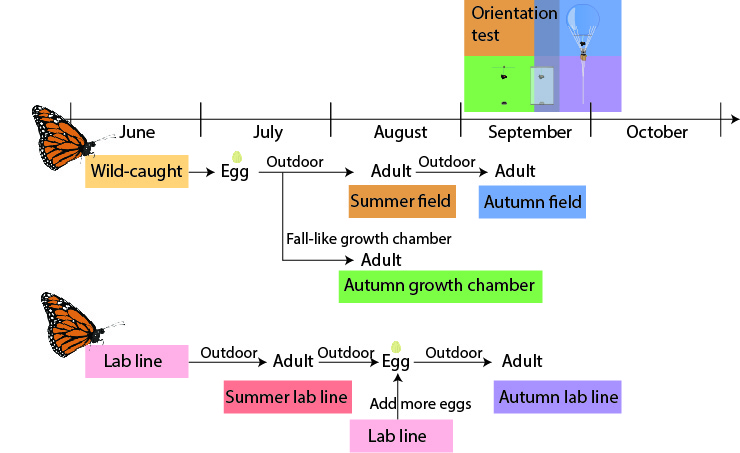

### Supplementary Figure 2

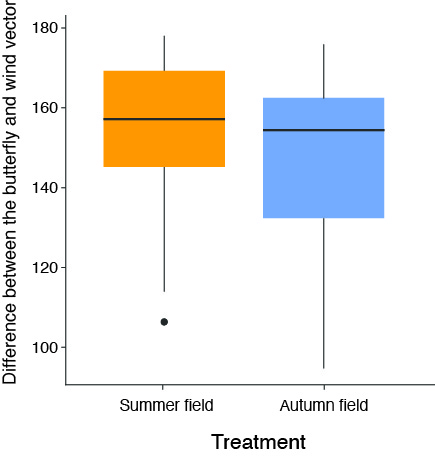

### Supplementary Figure 3

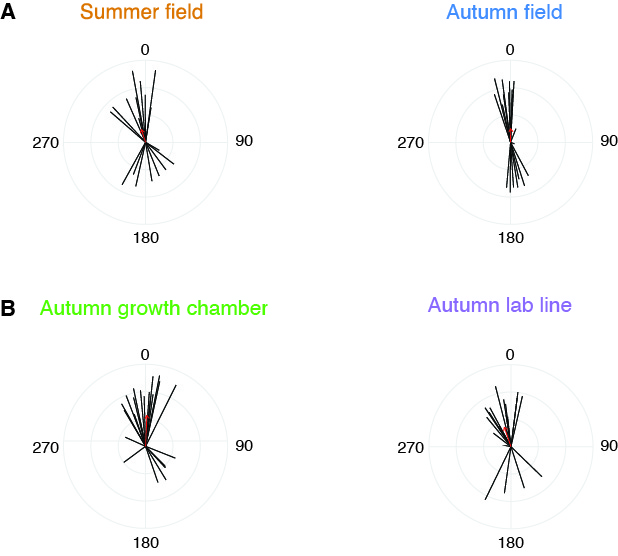

### Supplementary Figure 4

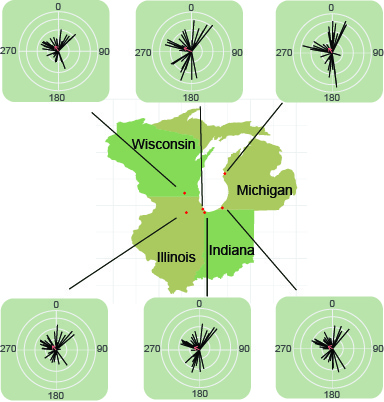
